# Partitioning phenotypic variance due to parent-of-origin effects using genomic relatedness matrices

**DOI:** 10.1101/133827

**Authors:** Charles Laurin, Gabriel Cuellar Partida, Gibran Hemani, George Davey Smith, Jian Yang, David M Evans

**Affiliations:** MRC Integrative Epidemiology Unit, University of Bristol, United Kingdom; University of Queensland Diamantina Institute, Translational Research Institute, Brisbane, Queensland, Australia; Institute for Molecular Bioscience, Queensland Brain Institute, The University of Queensland, Brisbane, Queensland, Australia

## Introduction

Parent-of-origin effects (POEs) describe the phenomenon in which the effects of alleles depend upon their parental origin. POEs imply that heterozygote individuals have phenotypes which are distributed differently depending upon which of their alleles were maternally and paternally transmitted (Guilmatre and Sharp 2012; Lawson et al. 2013). The extreme case of POEs is *polar overdominance*, where the two heterozygotes' phenotypes differ in distribution but the two homozygotes share the same distribution (Hoggart et al. 2014). Imprinting, a phenomenon in which one parent's allele is not expressed, is probably the most widely studied example of POE (Peters 2014).

POEs have traditionally been examined in the context of development, where, in mouse models, they have been associated with body size and social behavior (Peters 2014). One evolutionary explanation of POEs concerns genomic conflict between maternal and paternal genes in offspring, with paternal genes encouraging growth and solicitation of maternal care, even at the expense of the mother's health, while maternal alleles are orientated toward success of all offspring, which do not necessarily share paternity (Patten et al. 2014). Other evolutionary explanations for POEs include: different territorial patterns in males and females and coadaptation of maternal and offspring genomes to maximize the efficiency of nurturing behaviors like suckling and grooming (Peters 2014).

Whilst there is considerable support for the importance of POEs in animals (Neugebauer et al. 2010; Lawson et al. 2013), evidence for the existence of POEs in the etiology of complex human traits and diseases is mixed, in part due to the relative paucity of genomic data from families (Kong et al. 2009; Guilmatre and Sharp 2012). Before the genomics era, the children-of-twins design (Nance and Corey 1976), pedigree analyses (Hall 1990), and parent-offspring regressions (Clemons 2000; Smith et al. 2007) provided some limited evidence for the existence of POEs in human populations. These approaches were not often able to distinguish parental effects (indirect effects of the parental genotype on offspring phenotype) from POEs (interaction between the sex of the transmitting parent and the direct allelic effect in offspring) (Hager et al. 2008). More recently, in the genomics era, genome-wide association studies incorporating parent-of-origin information have been used to identify POEs at individual loci for age at menarche (Perry et al. 2014) Type I diabetes (Wallace et al. 2010), Type II diabetes, basal cell carcinoma, and breast cancer (Kong et al. 2009).

We propose a method, *G-REMLadp*, to estimate the phenotypic variance due to POEs across the genome by applying restricted maximum likelihood (REML) to offspring genome-wide genetic relatedness matrices and phenotype data. Our method involves the construction of a genetic relationship matrix indexing the parental origin of offspring alleles using genome-wide SNP data from parent-child duos or trios. The proposed method is an early adaptation to human genetics of procedures developed for animal breeding (Schaeffer et al. 1989; Spencer 2002; Nishio and Satoh 2015). The genotypic coding for POEs, which we adopt here, is based on the model outlined by Spencer (2002, 2009) and implemented by Nishio and Satoh (2015) in the context of genomic prediction. The genotype coding we used for dominance effects is due to Zhu *et al.*, (2015), who designed it to be orthogonal to the allele count at a locus; it is also orthogonal to the POE coding, which distinguishes our approach from that used by Spencer, Nishio and Satoh, and others. Using these codings, the phased genotype at a locus is coded using three orthogonal components: an additive-coded genotype, a dominance-coded genotype, and a POE-coded genotype.

To estimate the power of *G-REMLadp* to estimate non-null POEs, we provide an approximation using Haseman-Elston regression (Elston et al. 2000; Chen 2014). We also used simulated data to estimate the power and Type I Error rates of *G-REMLadp*, as well as the sensitivity of its variance component estimates to violations of assumptions. We then applied *G-REMLadp* to 38 phenotypes related to body size and obesity, metabolic traits, and IQ in a sample of up to 4689 individuals from a UK-based longitudinal study of childhood health and development.

## Methods

*G-REMLadp* uses REML to fit a linear mixed model incorporating random additive effects, dominance effects, and POEs to human genomic data, with the goal of partitioning phenotypic variance into components reflecting these sources of variation tagged by genome-wide SNP chips. In this model, at each locus, the phased genotype of each individual (*X* ∈ *aa*, *a_mo_ A_fa_*, *A_mo_ a_fa_*, *AA*) is expressed as 3 orthogonal components, which are then standardized. The codings for each of the 3 components are listed in Table 1. The standardized additive-coded genotype is the standardized minor allele (A) count at the locus. The derivation of the standardized dominance-coded genotype is given in Zhu *et al.* (2015); and a flexible equivalent coding is given in Álvarez-Castro (2015). The POE-coded genotype is −1 for the paternal-minor heterozygote (*a_mo_ A_fa_*), where *a_mo_* indicates that the major allele was inherited from the mother and *A_fa_* that the minor allele was inherited from the father), as 1 for the maternal-minor heterozygote (*A_mo_ a_fa_*), and 0 for both homozygotes, and is then standardized.

**Table I:** Recoding a phased genotype using three orthogonal terms.

| Phased Genotype | Freq | Add Code | Std Add | Dom Code | Std Dom | POE Code | Std POE |
| --- | --- | --- | --- | --- | --- | --- | --- |
| $a_{mo}a_{fa}$ | $q^2$ | 0 | $-\sqrt{2p/q}$ | 0 | $-p/q$ | 0 | 0 |
| $a_{mo}A_{fa}$ | $pq$ | 1 | $(q-p)/\sqrt{2pq}$ | $2p$ | 1 | -1 | $-1/\sqrt{2pq}$ |
| $A_{mo}a_{fa}$ | $pq$ | 1 | $(q-p)/\sqrt{2pq}$ | $2p$ | 1 | 1 | $1/\sqrt{2pq}$ |
| $A_{mo}A_{fa}$ | $p^2$ | 2 | $\sqrt{2q/p}$ | $4p-2$ | $-q/p$ | 0 | 0 |

*The minor allele is A, with frequency p, major allele a with frequency q = 1 - p. $A_{mo}$ : maternally inherited A allele; $A_{fa}$ : paternally inherited A allele. "Freq": expected frequency of the genotype under Hardy-Weinberg equilibrium. "Add Code": additive coding of the phased genotype. "Std Add": standardized additive coding based on mean 2p and variance 2pq. "Dom Code": dominance coding of the phased genotype. "Std Dom": standardized dominance coding based on mean 2p<sup>2</sup> and variance 4p<sup>2</sup>q<sup>2</sup>. "POE Code": parent-of-origin effect coding of the phased genotype. "Std POE": standardized parent-of-origin effect coding based on mean 0 and variance 2pq.*

The mixed model is given in Equation 1, where **y** is a vector of phenotypes; **μ***_y_* is a column vector of means; the vector **Tb** includes any fixed effects of covariates **T**; the **Z**s are *n* × *m* matrices containing the standardised coded genotypes, represented as one individual per row (sample size *n*) and one SNP per column (m markers affect the phenotype; in empirical data, this will be replaced by *m_o_*, the number of observed markers, while in power analyses of empirical data, this will be replaced by *m_e_*, the number of effective markers); the **β** are *m*-vectors of additive effects, dominance effects and POEs, which are assumed to be independently normally distributed with mean 0 and variances that are inversely proportional to the number of markers affecting the phenotype 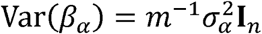 (**I** is the *n* × *n* identity matrix), Var (*β_△_*) = 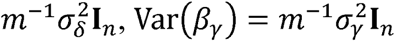, respectively, and **ε** is a normal error variable with mean 0 and variance 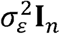 that is independent of the other variables on the right hand side of the equation. Figure 1 gives the phenotypic means at each phased genotype in the upper panel, while its lower panel illustrates how the phased genotype is represented as the three orthogonal standardized coded genotypes.

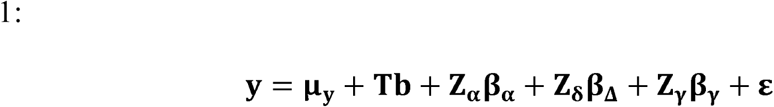

**Figure 1.**
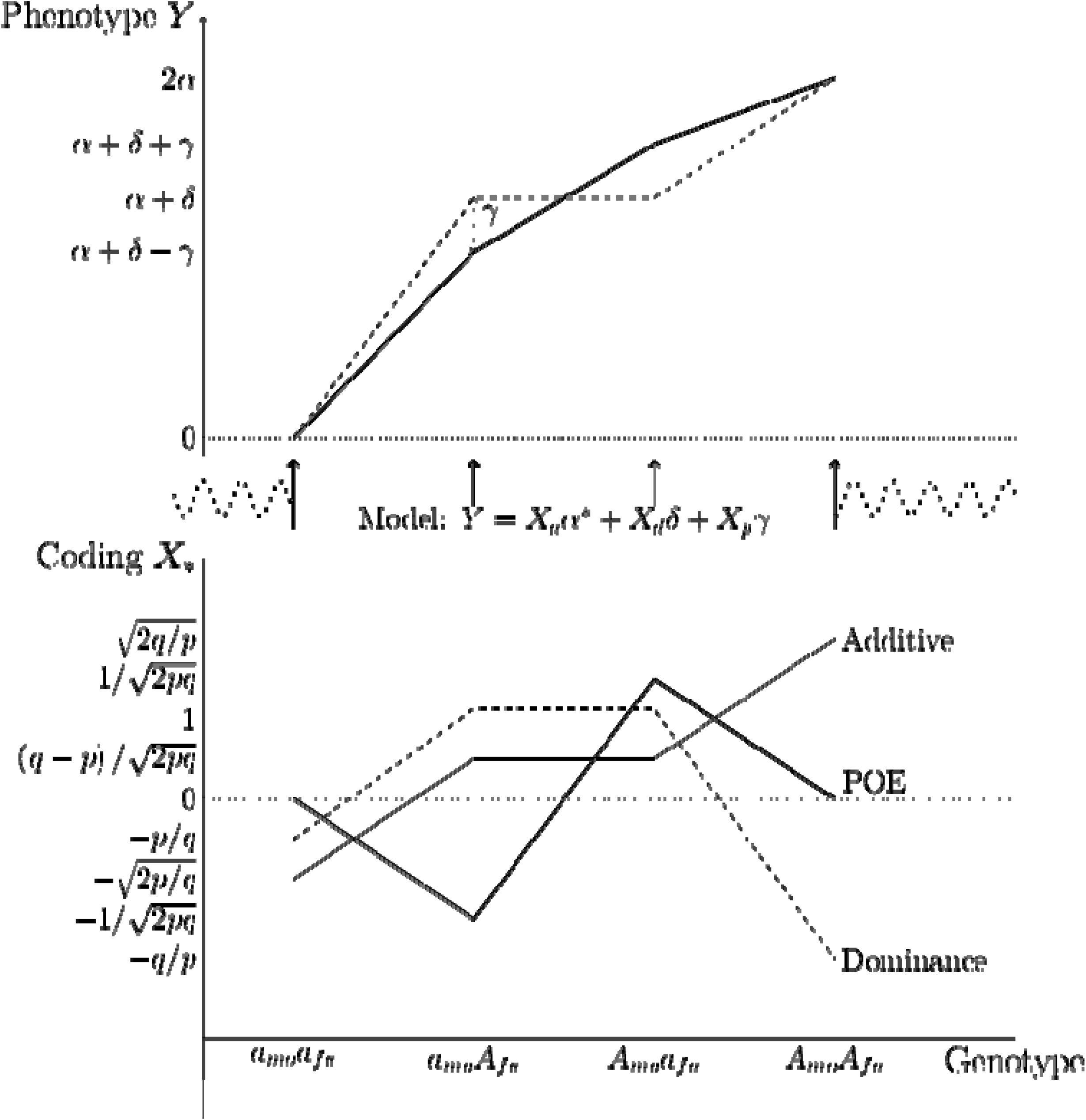
Phenotypic mean by phased genotype (top) Phased genotype represented by 3 standardised codings (bottom): (TOP)The dark line connects phenotypic means at different phased genotypes under a model including dominance and parent-of-origin effects in addition to additive effects. The dashed line connects phenotypic means under a model with additive and dominance effects only; the dominance deviation makes it so that the slope between the major-homozygote and the heterozygotes is steeper than that between the heterozygotes and the minor-homozygote (See Supplementary Figure S1). The difference due to parent-of-origin effects is illustrated by the dotted line marked *γ*. *a*: the major allele; *A*: minor (reference) allele; *a_mo_* : allele from mother; *a_fa_* : allele from father; *a*\* : additive allele-substitution effect; *δ* : dominance deviation; *α = α\** − (1 − 2*p*)*δ*, half the difference between homozygotes' phenotypic means; *γ*: the difference between the phenotypic mean of all heterozygotes and the phenotypic mean of paternal-minor heterozygotes. (BOTTOM) For a single locus with minor allele frequency *p*, the standardised additive coding (0,1,2 standardised to 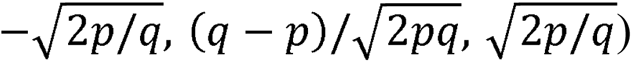 of the phased genotype is given by the medium-weight line. The standardised dominance coding is the dashed line (0,2*p*, 4*p* − 2 standardised to−*p*/*q*, 1, −*q*/*p*). The parent-of-origin effect coding is the thick line (0,−1,1,0 standardised to 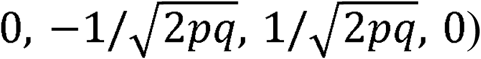.

The primary concern of this paper is with the partitioning of phenotypic variance components according to Equations 2 and 3, which present the problem in vector and individual-based forms, respectively.

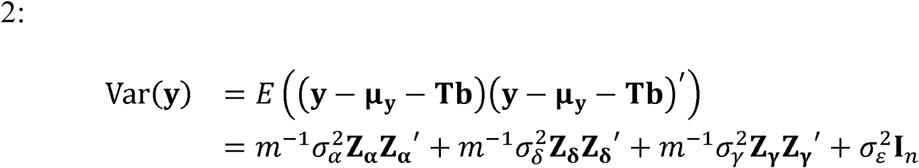

where the matrices **ZZ**′ are *n* × *n* matrices giving the cross-products, across all loci, of individuals' coded genotypes. The elements of Var (**y**) are given by

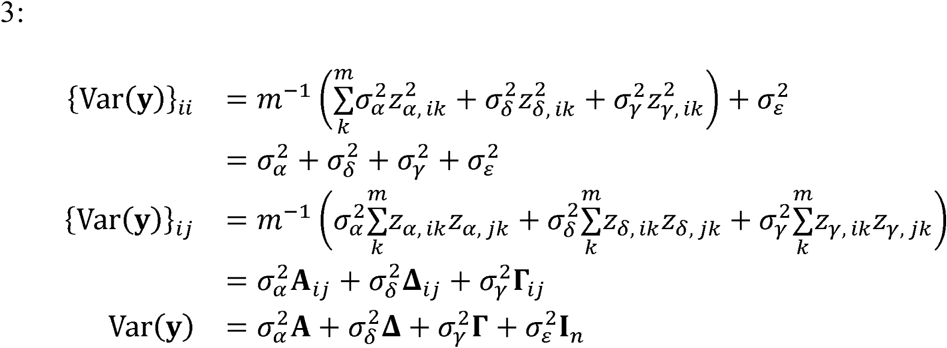

The second equality in Equation 3 is because the mean squares of each individual's standardized genotype codings are expected to be 1 in the absence of inbreeding.

The bold-faced Greek characters on the final line of Equation 3 are used to denote the genetic relationship matrices (GRMs) for the three coded genotypes; the additive GRM is, the dominance GRM **Δ**, and the POE-coded GRM is **Γ**. These are are averages over all effective markers of the sums of squares and cross-products of individuals' standardized coded genotypes.

## Statistical Methods

### Assumptions

Using REML to estimate the variance components model given in Equation 2 requires that several assumptions (both statistical and genetic) be met in order for inference about the model to be legitimate. In addition to the normality assumptions given above, *G-REMLadp* fitting of the model in Equation 3 assumes Hardy-Weinberg Equilibrium (HWE) and accuracy of phasing. Departures from HWE (such as non-random mating and differential allele frequencies in male and female parents (Falconer and Mackay 1996)) break the orthogonality of the POE-coded genotype with the additive-coded genotype and the dominance-coded genotype. Non-random mating could decrease the frequency of heterozygotes, while sex-specific allele frequencies cause the frequencies of the *a_mo_ A_fa_* and *A_mo_ a_fa_* heterozygotes to differ. Accurate phasing is required so that the different heterozygotes are correctly called; if they are not, *Zγ* is measured with error, diluting estimates of the *Zγ*, *y* association, hence downward-biasing estimates of 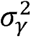.

### Power analysis via Haseman-Elston regression approximation

The power of G-REML analysis can be approximated in a Haseman-Elston (HE) regression framework where each distinct pair of individuals in the sample is used as the unit of analysis (Elston et al. 2000; Chen 2014; Visscher et al. 2014). In this framework, the outcome variable is the centered cross-product of phenotypes for each pair of unrelated individuals in the analysis (denoted *Y_ij_* for individuals *i* and *j*) and the predictor variables are the GRM entries under additive-coding (*A_ij_*), dominance-coding (*Δ_ij_*), and POE-coding (*Γ_ij_*) of the pair. The coefficients of the predictors in the associated ordinary least squares regression are estimates of the phenotypic variance due to each type of effect. This is similar to unweighted least squares estimation in covariance structure modelling (Browne 1982). Wald tests of significance for the variance components can be made using the assumption that the residuals in the regression are approximately normally distributed (Chen 2014).

The coded genotypes are orthogonal at each locus, meaning that the, **A**, **Δ** and **Γ** matrices are expected to be uncorrelated with increasing sample size and number of loci (assuming random mating). For example,

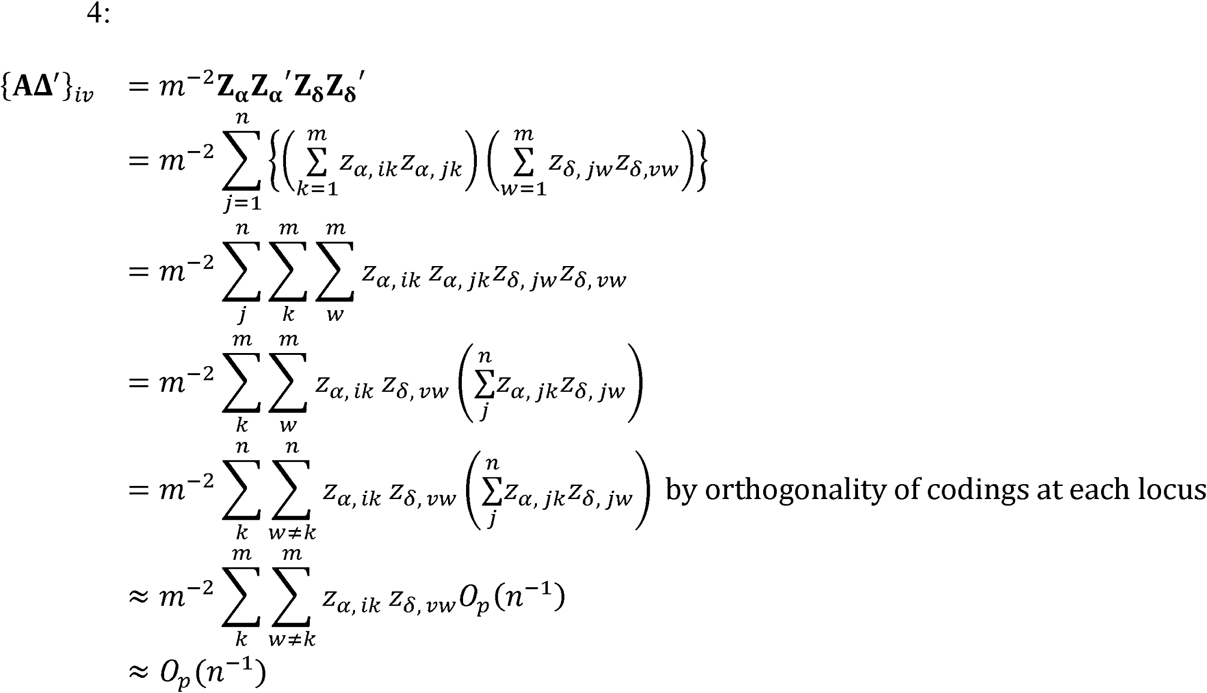

The sum over individuals *j* (in parentheses starting in the fourth line of the equation) will have mean 0 and variance *n*^−1^ over repeated sampling, which we denote using the *O_p_* notation to imply that its value is bounded in probability by Chebychev's inequality as *n* increases (Bishop et al. 1975, ch 14); the equality is approximate because the sum over individuals is not strictly uncorrelated with 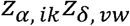. To get this wide bound, we also assume that the product of the additive and dominance codings between individuals *i* and *v* averages to be 1 over loci; because the product of the additive and dominance codings for two loci both with minor allele frequency *p* is 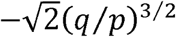, this assumption might be violated if many rare variants were used in the analysis. In the empirical analysis, the off-diagonal elements of the GRMs were uncorrelated 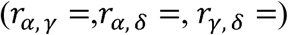; Supplementary Figure S2 illustrates this; while Supplementary Table S3 shows that the diagonal elements of the empirical GRMs were close to 1.

Equation 4 means that simple HE regressions of Y on each coded genotype's GRM will yield the same estimates (and associated Wald test statistics) as a multiple regression, and we can analyze the power of simple regressions but fit multiple components in practice. The Wald test statistic of a variance component, POE for example, is given by 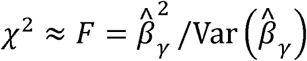 = 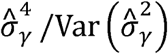. By assuming that a given variance component (i.e. the HE slope) is nonzero, the power of this Wald test depends on the noncentrality parameter of the associated statistic. The derivation of the noncentrality parameter is parallel for each type of coded genotype, so we focus on HE regression of POEs, following the example given for additive effects by Visscher *et al.* (2014).

Starting with standardized outcomes and predictor (where *vech* is the operator which transforms a symmetric matrix to a column vector of its lower-diagonal elements (Henderson and Searle 1979)). Simplifying assumptions are required: 1) *Y* is approximately normal (very unlikely for a cross-product phenotype) so that the Wald test statistic has an *F* distribution with 1 degree of freedom in the numerator and 1/2 *n*(*n* − 1) − 2 degrees of freedom in the denominator; and 2) the amount of variance in *Y* explained by *W* is so small and the sample size so large that the Wald *F* test statistic is well approximated by a 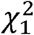 random variable.

We derive results for a set of *m* independent loci. The numerator of the Wald test statistic is the square of the estimated regression coefficient 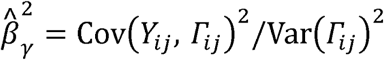. For individuals *i* and *j*, the expected covariance between *Y* and *Γ* is

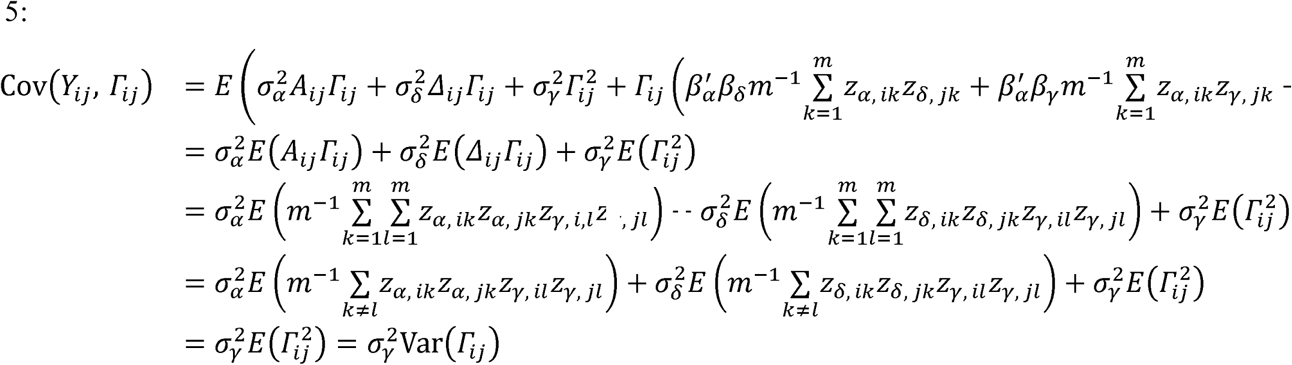

recalling that the coded genotypes are orthogonal within a locus. This derivation shows that the HE regression coefficient in the population is 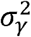, hence the numerator of the Wald test statistic is 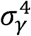.

The denominator of the Wald *F* test statistic is the error variance per degree of freedom, divided by the variance of the predictor variable, i.e. 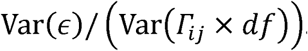. Here we use the approximation that Var (*ε*) ≈ Var(*Y*) = 1. HE regression uses the distinct pairs of observations as the units of analysis, so the denominator degrees of freedom are 1/2 *n*(*n*−1) when estimating the variance component and an intercept term. The variance of the POE GRM calculated at any single locus can be shown by direct calculation to be 1, which is also true for the additive effect and dominance effect GRMs (given assumptions). Var(*Γ_ij_*) is interpretable as the variance of the average of these component GRMs over *m_e_* loci, hence is 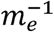 in this simple model. A more accurate approach (for additive effects) is given in Appendix 1 of Goddard (Goddard 2009), in which the variance in relatedness is averaged over the number of effective loci given variation in pedigree as well as linkage disequilibrium. Visscher *et al.* give an empirical estimate of the number of effective loci for GRMs calculated using genome-wide common SNPs in the HapMap3 panel, hence they suggest using Var (*A_ij_*) = 2 × 10^−5^ (Visscher et al. 2014). In our model, the denominator of the Wald test statistic is 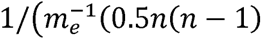 −2)) ≈ 2*m_e_* /*n*^2^. This means the mean Wald test statistic is approximately 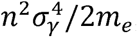, which is referred to a 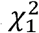 distribution because the denominator degrees of freedom in the *F*-test are very large. This derivation is nearly identical to that used in (Visscher et al. 2014), so it is possible to use their online tool (http://cnsgenomics.com/shiny/gctaPower/) to determine the power to detect a variance component of a particular size.

For *m_e_* loci and sample size *n*, the noncentrality parameters of the 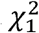 test statistic are:

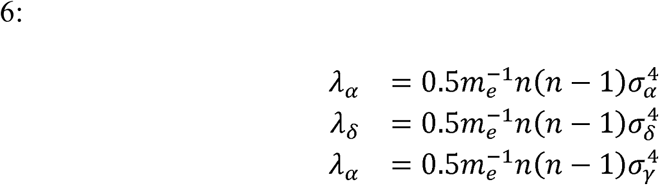

In our simulations, we used the true *m*, as loci were simulated without LD; in practice, we recommend the approximation 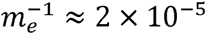 (Visscher et al. 2014), although its application to dominance effects and POEs is based only on analogy with additive effects and the number of effective loci will differ depending on which SNP panel is used.

### Implementation

*G-REMLadp* requires a set of phased genotypes, each with parent-of-origin assignments. Mitochondrial DNA and X chromosome SNPs are excluded from analysis. In the empirical analysis, we used a *Perl* script to assign parent-of-origin to genotypes which had already been phased, as described below. Given a set of phased, parental-origin-assigned SNPs, we first recoded each genotype to the three-term orthogonal coding given in Table 1, stored this data in the software package *MACH*'s (Li et al. 2010) “dosage” format, and then called *GCTA* in order to generate GRMs for the three variance components. *GCTA*'s --make-grm and --make-grmd (Zhu et al. 2015) make the appropriate GRMs for the additive and dominance effects directly from the *MACH*-formatted phased genotype. However, for POEs, we recoded each locus (in *R*) by subtracting the paternal minor allele indicator from the maternal indicator and writing this to an appropriately formatted .mldose.gz file, then generated an .mlinfo.gz file from the relevant sources for the SNP data set. We input the .mldose.gz and .mlinfo.gz files to *GCTA*'s MACH dosage function (--dosage-mach-gz) to generate the POE GRM. We then input the additive, dominance, and POE GRMs to *GCTA*, with the phenotype and covariate files, in order to fit the mixed model and estimate variance components. We chose to estimate the variance components without constraining them to be positive so that the null distribution of test statistics would not be a mixture (Visscher 2006). We also used the AI-REML algorithm instead of the Newton-Raphson or EM algorithms in order to fit the model quickly.

### Simulations

We simulated data according to the *G-REMLadp* model in order to: 1) evaluate the predicted statistical power using the Haseman-Elston regression approximation; 2) test the bias and variance of the method in response to violation of its assumptions; and 3) assess computational requirements. In simulation studies, data were simulated according to Equation 1, and variance components were estimated using both HE regression and *GCTA* software. The goals were to estimate the bias and variance of the variance component estimators, the agreement between HE and *GCTA* estimates, and the empirical power and Type I error rates of the HE test.

The design factors were: ssample size *n* = 1000, 2000, 4000, 5000, 7500, *m* = 500, 1000, 3000, 5000 SNPs; 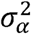/Var (y) = 0, 0.017, 0.033, 0.1;

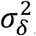/Var (y) = 0, 0.017, 0.033, 0.1; 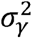/Var (y) = 0, 0.017, 0.033, 0.1; a violation of HWE—simulating maternal and paternal genotypes with different minor allele frequencies (either no difference, or mothers' allele frequency greater than fathers' by 0.05) so that the POE-coded genotype is no longer orthogonal to the other two codings (which remain mutually orthogonal); and a second violation of HWE-simulating parental genotypes to be correlated at *r* = 0 or *r* = 0.25, on average across all simulated loci, so that the diagonal entries of the GRMs are on average less than 1 and results might be less stable numerically. The sample size and number of loci used were constrained to be small due to computational requirements; computing GRMs at genome scale (*n* > 5000, *m_e_* > 10^6^) and fitting models to them required 100% use of at least 16 processors for approximately an hour, which is feasible for single empirical analyses but not in factorial simulations with thousands of replications. The SNPs were simulated to be independent with minor allele frequencies uniformly distributed between 0.01 and 0.5. Because there were relatively few of them, the simulated SNPs had much larger individual effects than would be expected in empirical data. We chose to simulate violations of HWE to test the robustness of *G-REMLadp* to violations of its assumptions, and as a way of widening the breadth of the simulations. Realistically, most samples in which *G-REMLadp* could be applied will have filtered SNPs for violations of HWE as part of routine QC (Laurie et al. 2010). A total of 7500 replications were simulated, but in the simulation results presented here, these factors were not completely crossed.

Three measures were used to evaluate the method: 1) empirical relative bias, the average difference between the simulated variance component and the value estimated in a replication, given as a percentage of the true value of the simulated variance component (except for null variance components, where we give the absolute bias); 2) empirical sampling variance, the variance of estimated variance components over replications; and 3) power/Type I error rate, the proportion of replications in which the Wald test of a null variance component exceeded the 95% critical value under a simulated non-null/null effect.

We expected power to increase with increasing samples size and variance component size and decreasing number of loci, and expected empirical variance to decrease according to the same pattern. We expected that violation of assumptions would lead to detectable bias.

### Empirical Analysis

We applied *G-REMLadp* to a sample of up to 4689 individuals gathered as part of the Avon Longitudinal Study of Parents and Children (ALSPAC), a prospective study of health and development beginning at pregnancy. We considered 38 different phenotypes which had been previously associated with POEs or which were related to body size, development, or social functioning.

See the Supplementary Material for further description of the sample, including the quality control procedures that we applied to it. The sample, the longitudinal study and its context, as well as its genotyping, are described in detail in two papers (Boyd et al. 2012; Fraser et al. 2013). The 38 phenotypes, along with estimates of additive effects, dominance effects, and POEs, are listed in Supplementary Table S1. All phenotypes were inverse-normal transformed prior to estimating variance components (Peng et al. 2007); fixed effects of sex and the first four ancestry-informative principal components were modelled. Parental effects and POEs have been previously studied in trios in this sample (Smith et al. 2007), but not genome-wide for the 38 phenotypes.

## Results

### Simulation Results

#### Power and sample size

Sample size curves at a Type I error rate of 5%, based on the HE approximation, are given in Figure 2, for POEs responsible for 1%, 3%, 10%, and 15% of the total phenotypic variance explained, tagged by 100000 loci. Figure 2 shows that a sample size of over 10000 genotyped duos or trios, with probands having phased genotypes, is likely necessary to detect the largest conceivable parent-of-origin effect variances and that a sample of 50000 individuals will be needed to detect POEs accounting for ≈ 1 – 3% of phenotypic variance, which was the size observed by Lopes *et al.* (2015). Figure 3 gives the empirical power, which was high because such large effects were simulated and because *m_e_* was low, with a median value of 2000 effective loci.

**Figure 2.**
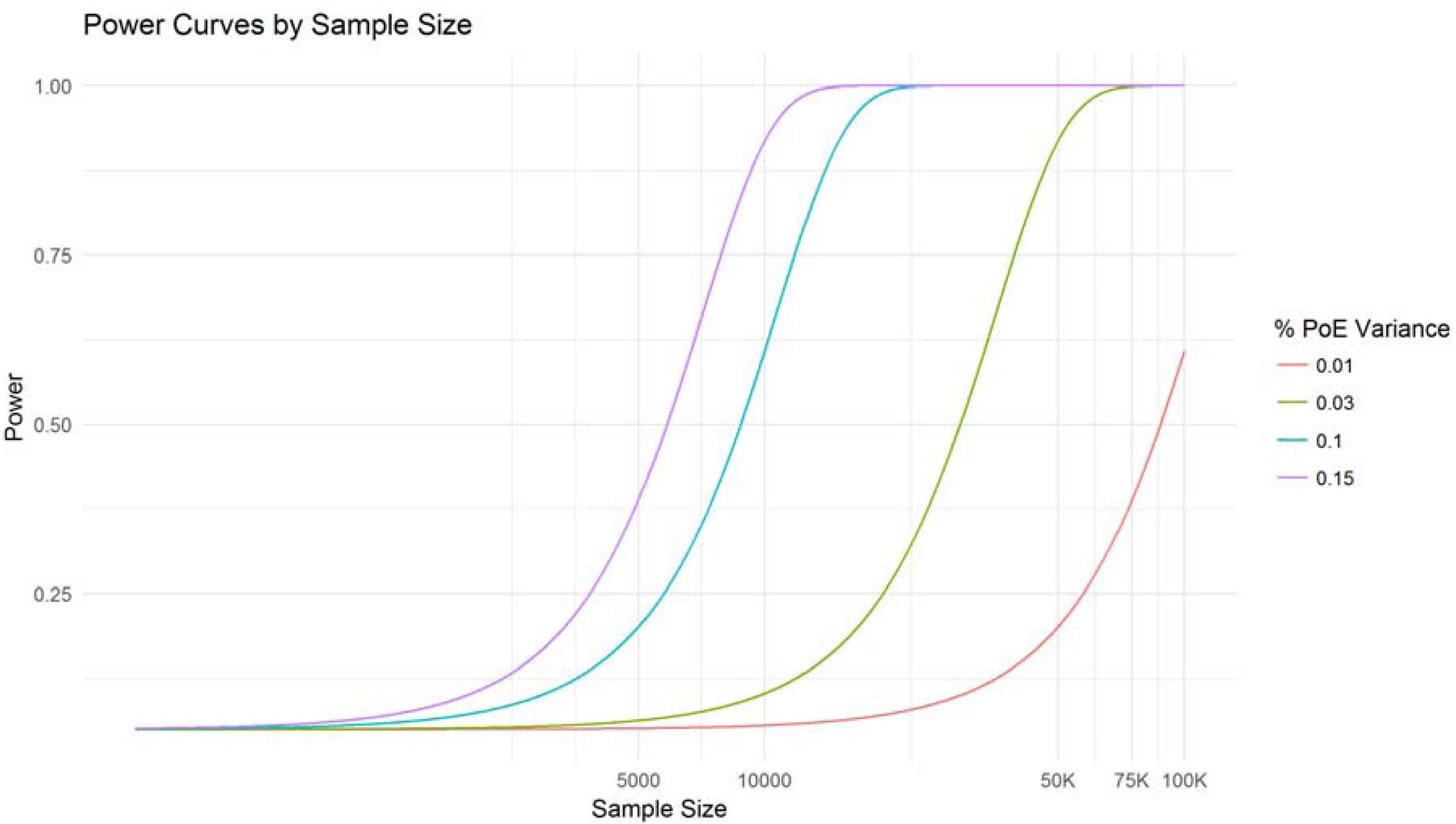
Expected power to detect parent-of-origin effects tagged by *m* = 10000 loci, by sample size: Curves generated using Haseman-Elston regression approximation to the Wald test of nonzero effect. %PoE: proportion of phentoypic variance attributable to parent-of-origin effects; Sample size: number of unrelated individuals with phased genotypes in the sample that would be used to estimate POEs; Power: proportion of test statistics generated under a non-null distribution with noncentrality parameter as given in Equation 4 that would exceed a 95% critical value in the Wald test of a null variance component.

**Figure 3.**
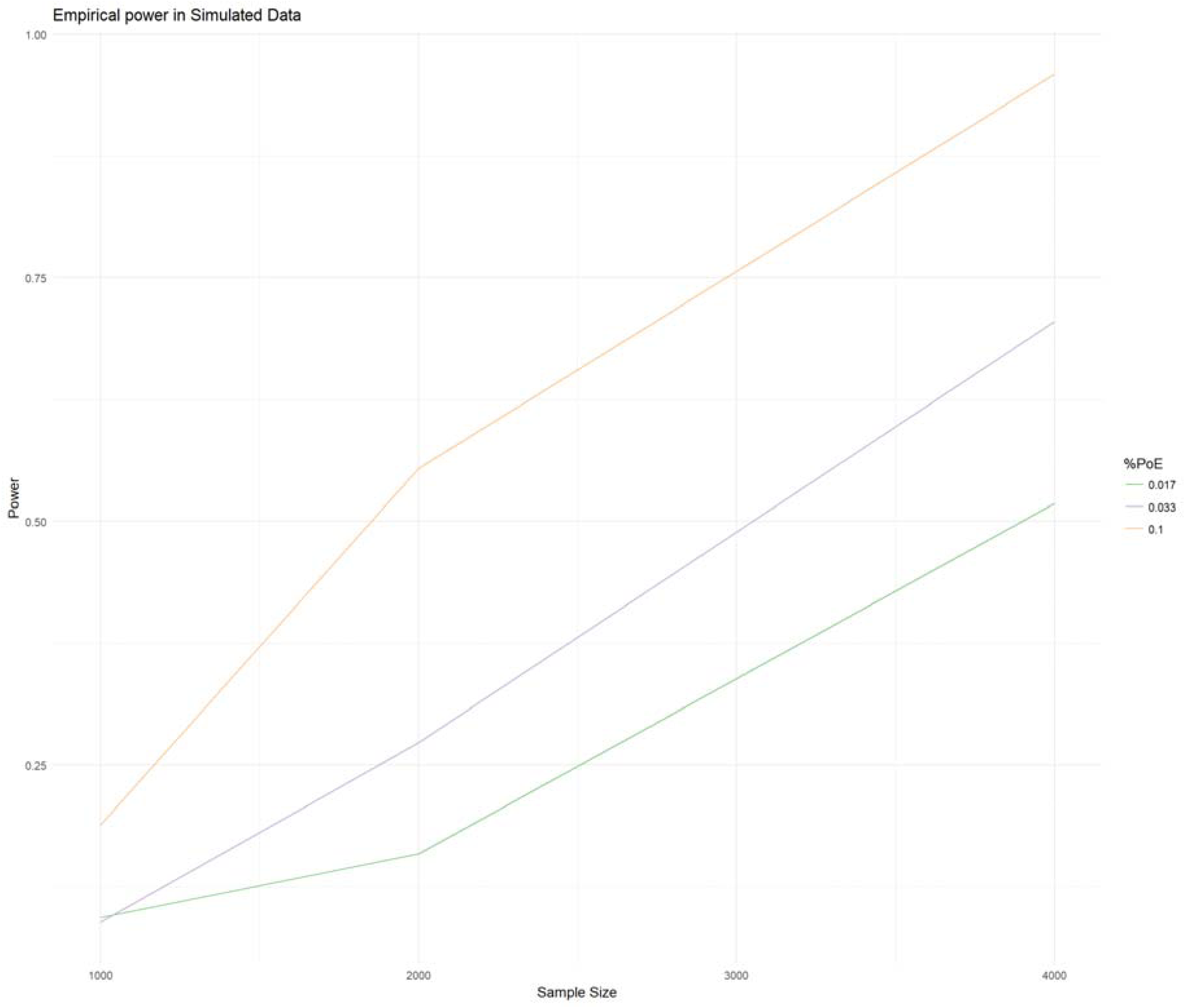
Empirical power curves in simulated data: Curves generated using proportions of significant Wald tests in simulated data with all assumptions specified in the text satisfied. %PoE: proportion of phentoypic variance attributable to parent-of-origin effects in the model used to generate simulated data; Sample size: number of simulated individuals with parent-of-origin determined at all loci; Power: proportion of replicates with Wald tests having *p*-values < 0.05. Note that the number of effective loci in this figure is an order of magnitude lower than in Figure 1.

The test involving HE regression is based on further simplifying and adding assumptions to the *G-REMLadp* procedure, hence discrepancies between the two approaches were expected, with the concern that large discrepancies would render the approximate power calculations unhelpful. Table S2, in the Supplementary Material, shows that the simulated HE-based test statistics lay relatively close to their expected values. Figures S1-S3 illustrate this graphically.

#### Type I error rates

There was no evidence that the tests of the POEs had inflated Type I error rates when assumptions were met; under these conditions, the simulated type I error rate was 0.0510. The HE-based test of the POE effect had slightly elevated Type I error rates (0.0625) under violation of HWE due to dependence of parental genotypes.

#### Bias and variance due to violating assumptions

We observed that violating HWE increased the discrepancy between predicted and observed test statistics. However, these results do not indicate whether this was due to increased bias or variance of the estimates, or both. Additionally, because the predicted test statistics were based on many simplifying assumptions, it is possible that the increased discrepancy would not have led to incorrect inference, and it is worth exploring whether violating HWE causes the estimated variance components to be misleading in predictable ways.

Results for simulations with truly null variance components are not shown. The absolute bias was under 3 × 10^−3^ for each type of effect whether or not assumptions were violated. Variances were similar to those for non-null estimates.

Table 2 shows the GCTA results under HWE. Relative bias was under 5% for additive, dominance, and parent-of-origin effects. Results were similar for HE model fitting.

**Table II:**
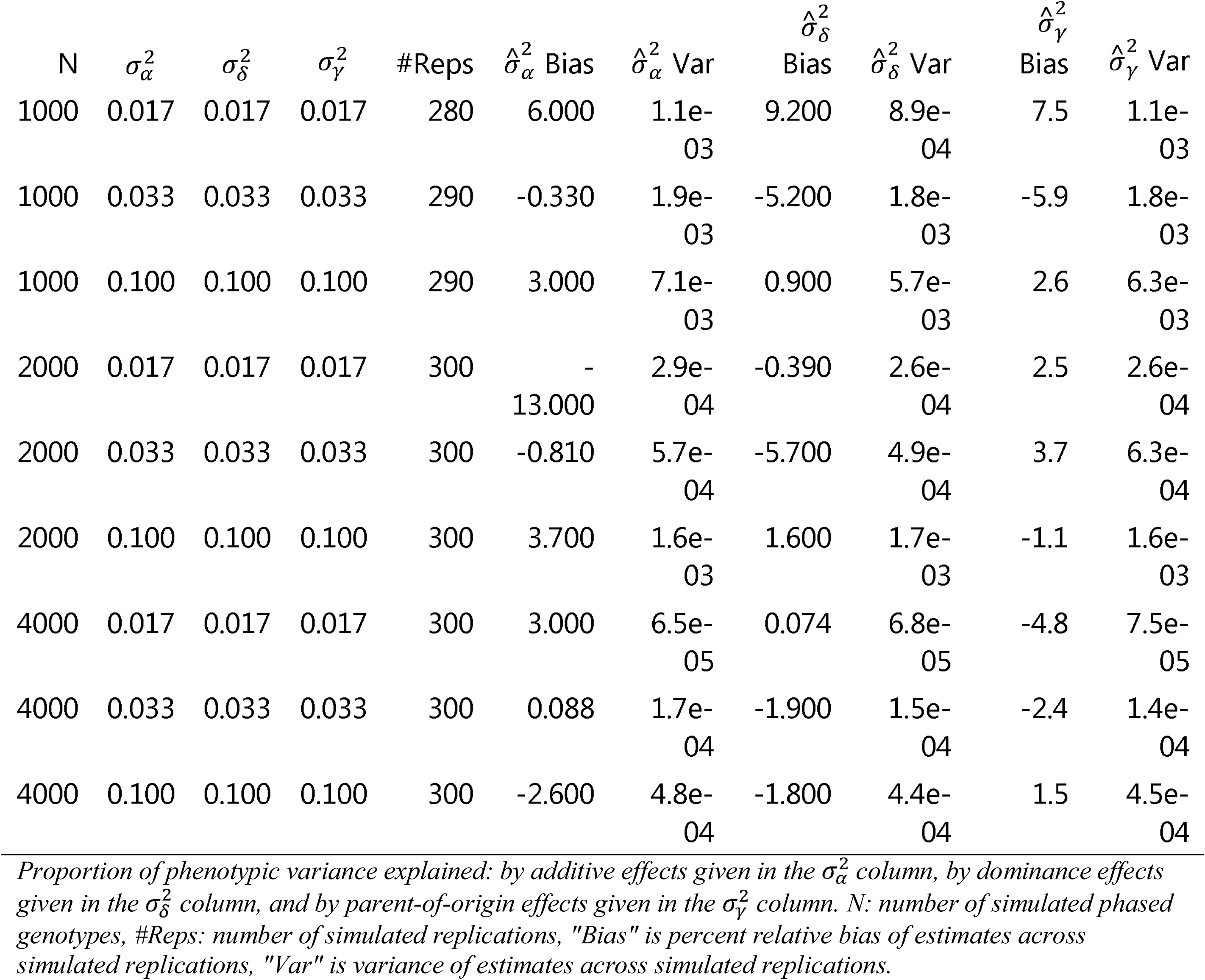
Relative bias and variance for G-REMLadp variance components when HWE was met, for non-null simulated effects.

Table 3 shows the GCTA results when the HWE assumptions were violated. Relative bias was under 10% for the additive and dominance effects but, when parental genotypes were correlated was downward biased by about 20% for the parent-of-origin effects. Results were similar for HE model fitting. Biases in estimating dominance effects were smaller but were uniformly positive.

**Table III:** Relative bias and variance for G-REMLadp variance components when HWE was violated, for non-null simulated effects.

| Parent Corr | MAF Diff | N | $\sigma_a^2$ | $\sigma_d^2$ | $\sigma_v^2$ | #Reps | $\hat{\sigma}_a^2$ Bias | $\hat{\sigma}_a^2$ Var | $\hat{\sigma}_d^2$ Bias | $\hat{\sigma}_d^2$ Var | $\hat{\sigma}_v^2$ Bias | $\hat{\sigma}_v^2$ Var |
| --- | --- | --- | --- | --- | --- | --- | --- | --- | --- | --- | --- | --- |
| 0.00 | 0.05 | 1000 | 0.017 | 0.017 | 0.017 | 300 | 10.00 | 1.0e-03 | -2.90 | 9.7e-04 | 17.00 | 1.0e-03 |
| 0.00 | 0.05 | 1000 | 0.033 | 0.033 | 0.033 | 300 | 3.20 | 2.4e-03 | 3.20 | 2.4e-03 | 2.00 | 2.0e-03 |
| 0.00 | 0.05 | 1000 | 0.100 | 0.100 | 0.100 | 300 | -1.00 | 6.6e-03 | 0.29 | 6.4e-03 | 0.31 | 5.7e-03 |
| 0.00 | 0.05 | 2000 | 0.017 | 0.017 | 0.017 | 300 | -8.40 | 2.5e-04 | 2.10 | 3.1e-04 | -0.11 | 2.8e-04 |
| 0.00 | 0.05 | 2000 | 0.033 | 0.033 | 0.033 | 300 | 2.80 | 5.4e-04 | 4.70 | 5.2e-04 | 0.90 | 4.8e-04 |
| 0.00 | 0.05 | 2000 | 0.100 | 0.100 | 0.100 | 300 | 2.20 | 1.6e-03 | -0.52 | 1.6e-03 | 0.69 | 1.6e-03 |
| 0.00 | 0.05 | 4000 | 0.017 | 0.017 | 0.017 | 300 | 1.10 | 7.2e-05 | -4.00 | 6.9e-05 | -5.60 | 8.1e-05 |
| 0.00 | 0.05 | 4000 | 0.033 | 0.033 | 0.033 | 300 | -1.90 | 1.5e-04 | 2.50 | 1.8e-04 | -2.50 | 1.5e-04 |
| 0.00 | 0.05 | 4000 | 0.100 | 0.100 | 0.100 | 290 | -2.10 | 4.9e-04 | 1.70 | 5.6e-04 | -0.20 | 4.3e-04 |
| 0.25 | 0.00 | 1000 | 0.017 | 0.017 | 0.017 | 300 | 3.60 | 8.2e-04 | 9.10 | 1.6e-04 | -8.40 | 1.1e-03 |
| 0.25 | 0.00 | 1000 | 0.033 | 0.033 | 0.033 | 290 | 5.70 | 1.6e-03 | 5.80 | 2.5e-04 | -20.00 | 2.0e-03 |
| 0.25 | 0.00 | 1000 | 0.100 | 0.100 | 0.100 | 290 | 1.70 | 5.6e-03 | 2.10 | 9.8e-04 | -25.00 | 8.3e-03 |
| 0.25 | 0.00 | 2000 | 0.017 | 0.017 | 0.017 | 300 | -0.80 | 2.0e-04 | -2.10 | 5.6e-05 | -33.00 | 2.8e-04 |
| 0.25 | 0.00 | 2000 | 0.033 | 0.033 | 0.033 | 300 | 6.00 | 3.9e-04 | -0.27 | 1.0e-04 | -21.00 | 6.5e-04 |
| 0.25 | 0.00 | 2000 | 0.100 | 0.100 | 0.100 | 300 | 3.60 | 1.5e-03 | 4.50 | 3.8e-04 | -25.00 | 2.2e-03 |
| 0.25 | 0.00 | 4000 | 0.033 | 0.033 | 0.033 | 300 | 0.68 | 1.2e-04 | 1.20 | 4.6e-05 | -21.00 | 1.3e-04 |
| 0.25 | 0.00 | 4000 | 0.100 | 0.100 | 0.100 | 300 | 1.30 | 3.6e-04 | 1.70 | 1.4e-04 | -23.00 | 4.7e-04 |
| 0.25 | 0.05 | 1000 | 0.017 | 0.017 | 0.017 | 130 | -24.00 | 8.4e-04 | 21.00 | 3.6e-04 | -21.00 | 1.3e-03 |
| 0.25 | 0.05 | 1000 | 0.033 | 0.033 | 0.033 | 160 | -3.50 | 1.7e-03 | 0.64 | 5.6e-04 | -45.00 | 2.5e-03 |
| 0.25 | 0.05 | 1000 | 0.100 | 0.100 | 0.100 | 220 | -0.91 | 5.2e-03 | 7.10 | 2.6e-03 | -31.00 | 8.2e-03 |
| 0.25 | 0.05 | 2000 | 0.017 | 0.017 | 0.017 | 280 | -5.70 | 1.9e-04 | 0.26 | 1.1e-04 | -28.00 | 3.2e-04 |
| 0.25 | 0.05 | 2000 | 0.033 | 0.033 | 0.033 | 300 | 0.25 | 3.8e-04 | 6.00 | 2.8e-04 | -26.00 | 5.4e-04 |
| 0.25 | 0.05 | 2000 | 0.100 | 0.100 | 0.100 | 300 | 5.20 | 1.4e-03 | 1.20 | 7.4e-04 | -21.00 | 1.8e-03 |
| 0.25 | 0.05 | 4000 | 0.033 | 0.033 | 0.033 | 300 | -0.87 | 1.3e-04 | 4.40 | 7.8e-05 | -23.00 | 1.6e-04 |
| 0.25 | 0.05 | 4000 | 0.100 | 0.100 | 0.100 | 300 | 2.20 | 3.1e-04 | 3.50 | 2.5e-04 | -23.00 | 4.8e-04 |

Parent Corr: average correlation of parental genotypes, MAF Diff: average difference in parental MAFs. Proportion of phenotypic variance explained by additive effects is given in the $\sigma_a^2$ column, by dominance effects in the $\sigma_d^2$ column, and by parent-of-origin effects in the $\sigma_v^2$ column. N: number of simulated phased genotypes, #Reps: number of simulated replications, "Bias" is percent relative bias of estimates across simulated replications, "Var" is variance of estimates across simulated replications.

The *GCTA* and HE estimates tended to differ in the presence of correlation between parental genotypes; in this situation, HE regression estimates of POEs had higher variance (and hence higher mean-squared error) than did *GCTA* estimates, which is expected to be the case even in additive-only models (Chen 2014). As a result, the correspondence between the two methods decreased from a HE-*GCTA* correlation of 0.9911(0.9908,0.9915) with independent parental genotypes to 0.9624(0.9597, 0.9649) with correlated parents. This was not observed for estimates of the two other types of effects.

#### Empirical Analysis

Estimates of 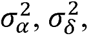 and 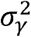 are given in Figure 4 for the 38 phenotypes. Error bars showing ± 1.96 standard errors are given for each estimate. Based on this, the sample size of n ≈ 5000 was sufficient to detect established additive effects for anthropometric variables such as height and fat mass. The mean estimate for additive effects was 0.297(0.10). There is evidence for dominance effects on verbal IQ and fat mass in the sample. The mean estimate for dominance effects was 0.031(0.14). The largest (and most reliably estimated) POEs were estimated for age at menarche, FVC, and age at first tooth. The mean estimate for POEs was 0.023(0.10).

**Figure 4.**
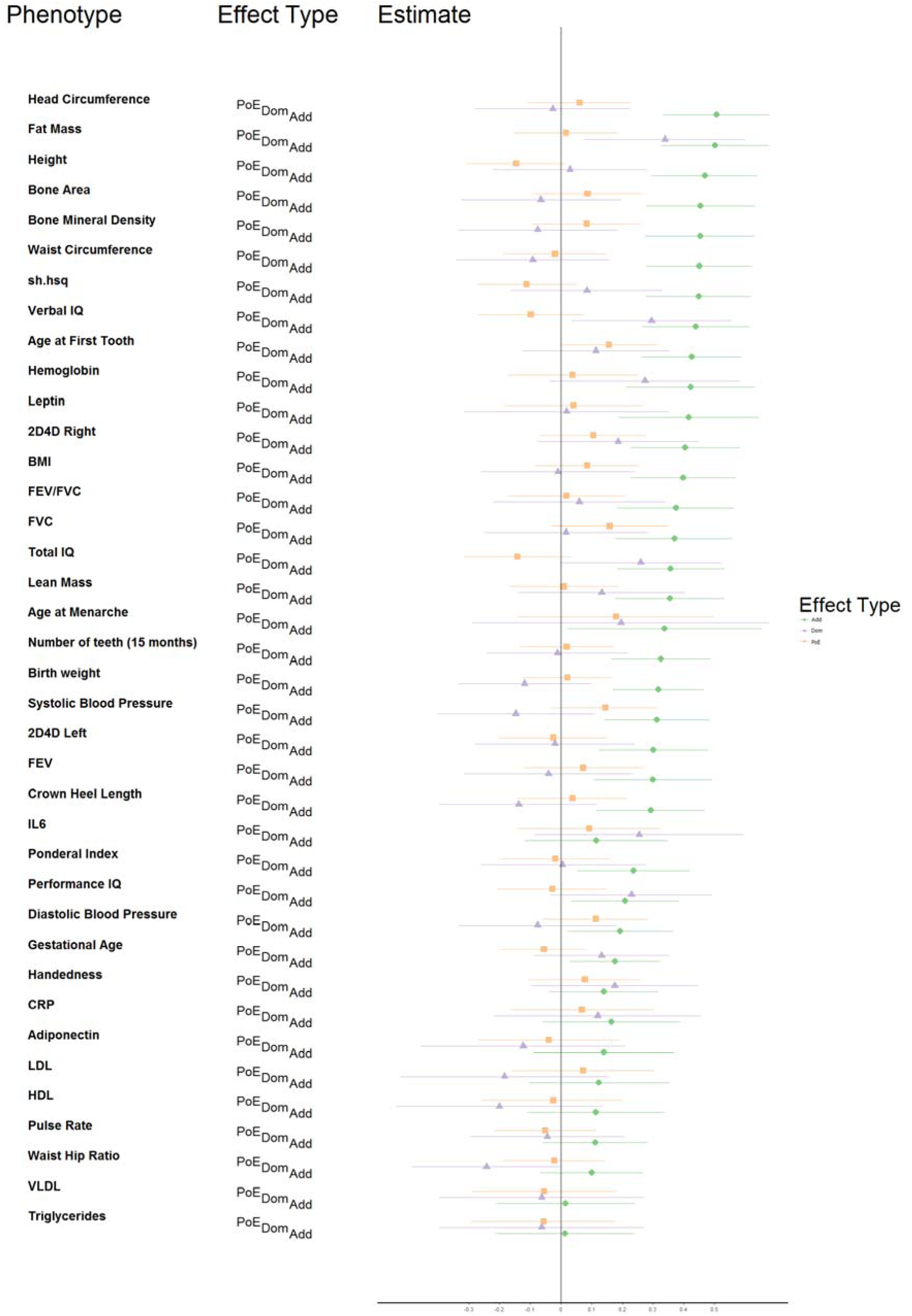
Forest plot of variance component estimates of 19 phenotypes in ALSPAC: The *x*-axis, labelled “Estimate,” is the proportion of phenotypic variance attributable to each effect type (PoE: parent-of-origin, squares; Dom: dominance, triangles; Add: additive, circles). To avoid bias, variance estimates were not constrained, hence negative variance component estimates are possible, for example if too many components are fit. Estimates are marked by the shapes, lying in 95% confidence intervals. A vertical line is plotted at 0.

Across the 38 phenotypes surveyed, which are listed in Table S1, in the aggregate, additive effects were frequently estimated reliably and away from 0. POEs had standard errors approximately equal to those of additive effects, while estimates of dominance variance showed slightly larger standard errors. This is illustrated using “dodged” (Wilkinson 2006) histograms of variance component estimates in Figure 5, which is broadly similar to that given (for additive and dominance effects) in Figure 2 of Zhu *et al.* 2015 (Zhu et al. 2015).

**Figure 5.**
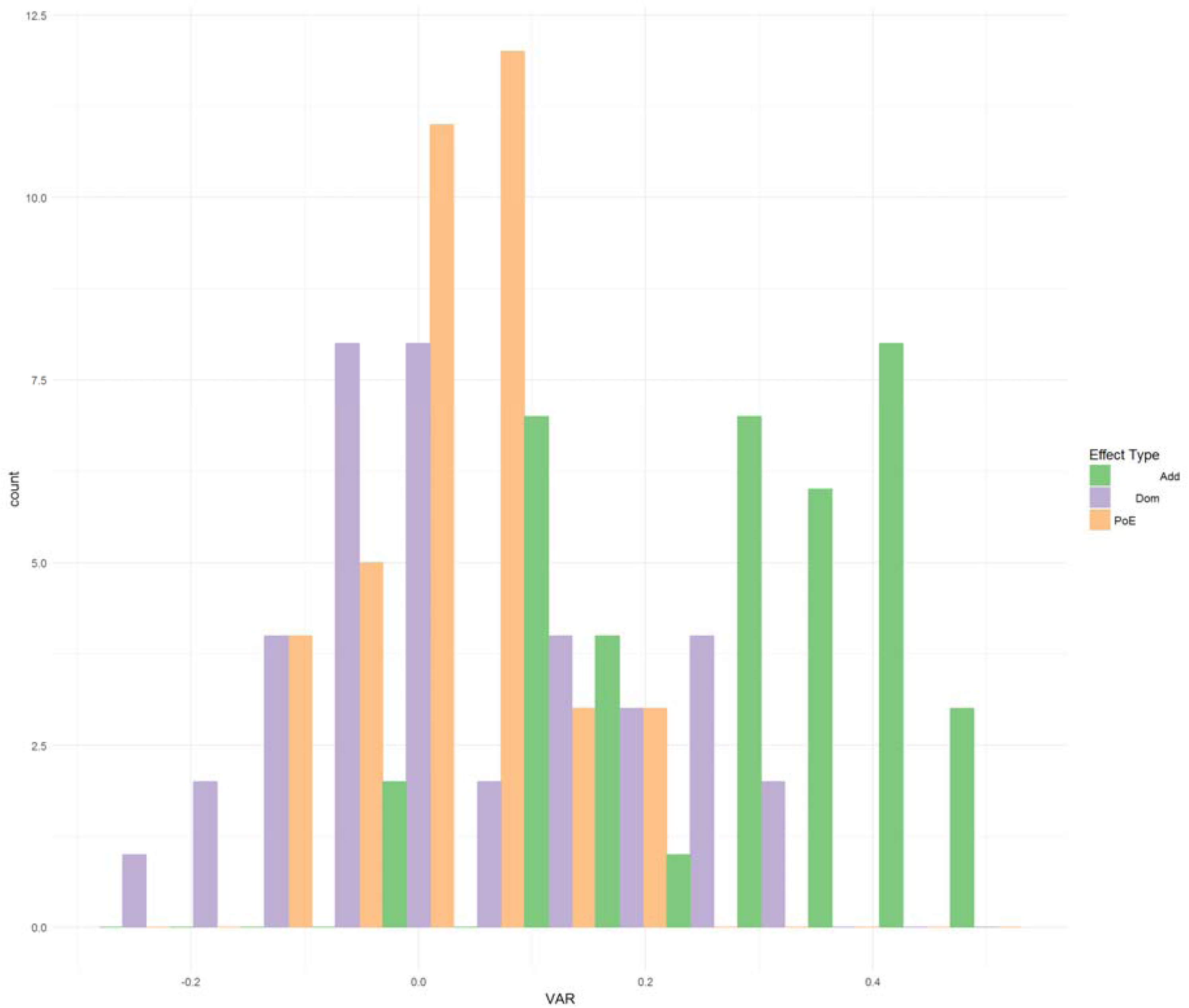
Histograms of the estimates of additive, dominance, and parent-of-origin variance components for 38 ALSPAC traits. Add: Estimated proportions of phenotypic variance attributable to additive effects; Dom: Estimated proportions of phenotypic variance attributable to dominance effects; PoE: Estimated proportions of phenotypic variance attributable to parent-of-origin effects. To avoid biased variance estimates, they were not constrained to be positive; hence, for POEs, which the study was underpowered to detect, there are many negative estimates.

Two patterns in variance partitions emerged (although the sample size is too small to perform significance tests of differences in heritability estimates across traits). Age at menarche, age at first tooth, CRP, handedness, IL-6, and 2d4d ratio in the right hand showed a pattern of relatively equal additive, dominance, and parent-of-origin effect variance estimates which were relatively far from 0 (though not usually greater than 2 standard errors from 0). A second pattern was seen in the IQ measures and fat mass, where additive and dominance effect estimates were approximately equal and nonzero. In the remaining 30 phenotypes, if an effect was detectable, it was an additive effect.

## Discussion

The most important findings from our simulations are that: 1) *G-REMLadp* does not seem to be inherently biased in estimating variance due to additive effects, dominance effects, and POEs, and 2) that substantial correlation between parental genotypes is necessary to bias *G-REMLadp* estimates. We did not investigate the effect that linkage disequilibrium might have on our results; if local levels of linkage disequilibrium are associated with POE size, an LDAK implementation of *G-REMLadp* may help address this (Speed et al. 2012). The empirical and simulation results agree that a sample size under 10000 is insufficient to generate precise *G-REMLadp* estimates. Hence, POEs of the size observed by Lopes *et al.* (Lopes et al. 2015) (≈ 2% of variance explained) are likely to require sample sizes close to 50000 in order to resolve properly (their samples were about 4500 purebred pigs, hence *m_e_* far below that for humans on the HapMap 3 panel, accordingly their pattern of results is closer to our simulated findings than to our empirical ones). Nevertheless, it may be interesting following up the ALSPAC results for age at menarche (a phenotype known to be influenced by imprinted loci (Perry et al. 2014)), age at first tooth, and 2d4d ratio results in a larger study with trios. Replicating the dominance results for fat mass would also be of interest, as Zhu *et al.* (2015) found no significant dominance for skinfold thickness; similarly, replicating a dominance heritability component of verbal IQ would substantiate venerable claims for non-additive effects on cognitive performance (Devlin et al. 1997; Plomin and Deary 2015).

However, the highest-profile claim (Devlin et al. 1997) for non-additive effects on IQ involves maternal effects. The evidence for this claim comes from findings of increased similarity in IQ in the children of female twins relative to the children of male twins (Nance and Corey 1976). Maternal (or paternal) effects are defined as indirect effects of the parent's genotype on offspring phenotype, usually via parental influence on offspring's environment (Wolf and Wade 2009). These are distinct from the POEs discussed here, because POEs are direct effects of the offspring's genotype on its phenotype. In the presence of maternal (paternal) effects, the mother's (father's) genotype explains some of the residual phenotypic variation remaining after regressing the offspring's phenotype on its genotype. However, methods designed to detect POEs by modelling different distributions of the two heterozygotes' phenotypes (including *G-REMLadp*) cannot distinguish POEs from certain patterns of maternal/paternal effects (Hager et al. 2008). It is also possible that there are maternal effects that cause imprinting (epigenetic changes due to intrauterine environment or family environment created by parental behavior). In order to distinguish POEs from maternal effects, one can fit a model with both maternal and POEs or restrict the analysis to offspring of heterozygous mothers. Given the sample size considerations in our results, the former approach is likely to be more successful; *G-REMLadp* could be implemented in OpenMx and used as an addition to an M-GCTA model (Eaves et al. 2014; Kirkpatrick and Neale 2015; Neale et al. 2015). However, POEs and maternal/paternal effects represent departure from Mendelian inheritance; a reliable nonzero 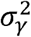 value detected by *G-REMLadp* is worthy of followup even if it represents a mixed bag of POEs and indirect effects.

Other than the insufficient sample size in the ALSPAC analyses, an additional limitation of this project was that the simulations and empirical data were not closely matched; less idealized, more informative simulations could have been performed. For example, simulated phenotypes could've been based on the empirical distributions of the ALSPAC phenotypes, and these, coupled with real GRMs (with imperfect HWE because of sampling error, if not actual violations), could've resulted in more thorough evaluation of *G-REMLadp*. Pairwise relatedness values could also have been simulated from the empirical distributions of (*A_ij_*, *Δ_ij_*, and *Γ_ij_*) or smooth approximations to them (Lee and Chow 2014). This approach is more feasible computationally under the HE regression framework than under maximum likelihood. *G-REMLadp* is not limited to genome-wide analyses; future work could also make use of relatedness at known imprinted loci, though this would require sufficiently large samples to account for the lower degree of tagging of phenotypic variation. A 2015 paper by Baran *et al.,* in *Genome Research*, supplies an atlas of such locations in the genome which affect expression (Baran et al. 2015). A related future direction (as suggested by Lopes *et al.*) would be to emphasize the connection between imprinting and attempts to detect POEs, for example by applying the method in the EWAS, rather than GWAS, context, i.e. identifying genome-wide eQTL effects on age at menarche and first tooth, and 2d4d ratio (Zhu et al. 2016).

In summary, *G-REMLadp* offers the ability to easily partition phenotypic variance according to three types of inherited effects. With large studies of parent-child trios/duos, it is possible to fit *G-REMLadp* models and detect effects without bias. Finally, failure to model POEs has been identified as one possible source of missing heritability (Kong et al. 2009; Eichler et al. 2010). For any phenotypes with missing heritability that is uncovered by modelling POEs, the epigenetic and evolutionary implications of POEs lead to hypotheses of distinctive etiologies and genomic architectures.

## Acknowledgements

This work was supported by NHMRC Project Grant (APP1085130 to D.M.E) and a Medical Research Council program grant (MC_UU_12013/4 to D.M.E). The UK Medical Research Council and the Wellcome Trust (Grant refs: 092731 and 102215/2/13/2) and the University of Bristol provide core support for ALSPAC. D.M.E is supported by an Australian Research Council Future Fellowship (FT130101709). J.Y. is supported by the Sylvia & Charles Viertel Charitable Foundation Senior Medical Research Fellowship. GWAS data was generated by Sample Logistics and Genotyping Facilities at the Wellcome Trust Sanger Institute and LabCorp (Laboratory Corporation of America) using support from 23andMe. We are extremely grateful to all the families who took part in this study, the midwives for their help in recruiting them, and the whole ALSPAC team, which includes interviewers, computer and laboratory technicians, clerical workers, research scientists, volunteers, managers, receptionists and nurses. This publication is the work of the authors and D.M.E will serve as guarantor for the contents of this paper.

